# Transcutaneous auricular vagus nerve stimulation during movement modulates motor neural circuitry without widespread cortical or autonomic activation

**DOI:** 10.1101/2025.07.07.663419

**Authors:** Cléo Perrin, Flaminia Pallotti, Tiziano Weilenmann, Clément Lhoste, Weronika Potok-Szybinska, Xue Zhang, Nicole Wenderoth, Olivier Lambercy, Dane Donegan, Paulius Viskaitis

**Affiliations:** Rehabilitation Engineering Laboratory, Department of Health Sciences and Technology, ETH Zurich, Zurich, Switzerland; Elite-Master of Science in Neuroengineering (MSNE), Technical University of Munich (TUM), Germany; Neural Control of Movement Laboratory, Department of Health Sciences and Technology, ETH Zurich, Zurich, Switzerland; Neuroscience Center Zurich (ZNZ), University of Zurich, Federal Institute of Technology Zurich, University and Balgrist Hospital Zurich, Zurich, Switzerland; Future Health Technologies, Singapore-ETH Centre, Campus for Research Excellence and Technological Enterprise (CREATE), Singapore, Singapore

**Keywords:** taVNS, EEG, TMS, context-paired, corticomotor excitation, CST excitability, pupillometry, arousal

## Abstract

Transcutaneous auricular vagus nerve stimulation (taVNS) is a promising neuromodulatory approach for treating neurological disorders, with growing interest in its potential to support motor rehabilitation. Yet, its mechanisms of action, potentially influenced by behavioral context, remain elusive. This sham-controlled study investigated transient taVNS interactions with movement in healthy adults, focusing on autonomic, neuromodulatory, and motor circuits. During a finger-tapping paradigm, heart rate (HR), galvanic skin response (GSR), pupil diameter, and electroencephalography (EEG) were recorded to probe movement-dependent stimulation effects. This study first identified a novel physiological dissociation: all measures responded to movement, but taVNS did not significantly alter HR, GSR, or general EEG spectral slope; taVNS increased pupil diameter in both conditions, but enhanced sensorimotor EEG spectral slope solely during movement. This context-specific effect on motor systems was further supported by a transcranial magnetic stimulation (TMS) experiment demonstrating increased corticospinal excitability during taVNS. These findings provide mechanistic insights into how taVNS may selectively enhance motor system responsiveness during active states, supporting future exploration of behaviorally paired stimulation protocols for neurorehabilitation.

## INTRODUCTION

Transcutaneous auricular vagus nerve stimulation (taVNS) is rapidly gaining traction as a non-invasive neuromodulation technique showing promising therapeutic effects (Butt et al. 2020; Ali et al. 2024; Frangos, Ellrich, and Komisaruk 2015) across diverse conditions such as stroke recovery (Yan, Qian, and Li 2022; Redgrave et al. 2018; Badran et al. 2023), epilepsy (von Wrede and Surges 2021), depression (C. Tan et al. 2023), and chronic pain (Costa et al. 2024). Yet, the understanding of how stimulating a single peripheral nerve can yield such broad therapeutic effects remains limited.

Anatomically, it is well established that the auricular branch of the vagus nerve cell bodies, located in the superior jugular ganglion, innervates the posterior wall of the external auditory meatus and cymba conchae, and projects to the nucleus tractus solitarius (NTS) in the brainstem (Butt et al. 2020; Ali et al. 2024; Frangos, Ellrich, and Komisaruk 2015; Kaniusas et al. 2019). In turn, the NTS recruits both descending autonomic pathways - including the parabrachial nuclei, periaqueductal gray, pontine raphe nuclei, and dorsal motor vagus nucleus (DMVN) (Zou et al. 2024; Toschi et al. 2023) - and ascending neuromodulatory pathways such as the noradrenergic, serotonergic, dopaminergic, and cholinergic systems (Ali et al. 2024; Yakunina, Kim, and Nam 2017; Frangos, Ellrich, and Komisaruk 2015; Borgmann et al. 2021). Subsequently, these systems broadcast signals across widespread brain networks, including those involved in attention, pain processing, and motor control; a pattern of recruitment that has been functionally confirmed by fMRI studies (Borgmann et al. 2021; Huang et al. 2023).

Importantly, neural systems targeted by taVNS are also recruited as part of the sensorimotor circuitry during movement, including ascending neuromodulatory pathways of noradrenergic and cholinergic systems (Collins et al. 2023), and autonomic processes (e.g. heart rate fluctuations during physical activity). This overlap suggests potential synergistic interactions between taVNS and movement, though the existence and nature of these interactions remain unknown. Yet, understanding the effects of taVNS during movement is clinically relevant, as transient taVNS has been paired with motor activity in neurorehabilitation with the goal of enhancing functional recovery (Yan, Qian, and Li 2022; Redgrave et al. 2018; Badran et al. 2023), supported by promising evidence of invasive VNS efficacy (Dawson et al. 2016; Kilgard et al. 2025). However, to date, most mechanistic studies of taVNS have focused on neurophysiological responses to taVNS in resting state paradigms (Clancy et al. 2014; Lloyd et al. 2023; Sharon, Fahoum, and Nir 2021; Wienke et al. 2023). This emphasis on resting states overlooks the inherently dynamic nature of neural processing during active behavior, particularly movement, when relevant networks are more robustly engaged (Zagha et al. 2022; Salkoff et al. 2020). As a result, a critical and largely unaddressed gap remains, which our study sought to answer: are taVNS effects during movement behavior the same as those at rest, and are there specific interactions between taVNS and movement within implicated neural pathways?

Therefore, in this study, we assessed neurophysiological markers associated with autonomic, neuromodulatory, arousal and sensorimotor pathways during a finger-tapping paradigm in healthy adults. We compared an active transient taVNS condition against a no stimulation control and an earlobe sham stimulation, across two movement-related states: an active (go) condition involving voluntary movement, and a motor inhibition (no-go) condition. More specifically, we assessed measures of heart rate (HR), galvanic skin response (GSR), pupillometry, and electroencephalography (EEG), where each modality indexes a distinct neurophysiological domain. HR and GSR reflect aspects of autonomic nervous system (ANS) activity, and are often used as indices of physiological arousal and metabolic readiness (Wang et al. 2018; Pop-Jordanova and Pop-Jordanov 2020). Pupil diameter serves as a sensitive and temporally precise proxy for ascending neuromodulatory systems - such as the locus coeruleus (LC) and lateral hypothalamus (LH) - and reflects fluctuations in arousal, attention, and internal brain states (Viglione, Mazziotti, and Pizzorusso 2023; Grujic et al. 2023; Pfeffer et al. 2022; Weijs et al. 2025). EEG, through spectral analysis (e.g., spectral slope), provides a measure of cortical excitability and balance of excitatory-inhibitory dynamics across distributed neural populations, enabling insight into state-dependent processing in sensorimotor and associative circuits (R. Gao, Peterson, and Voytek 2017; Weijs et al. 2025; Lendner et al. 2020). Finally, transcranial magnetic stimulation (TMS)-induced motor evoked potentials (MEPs), which provide a well-established index of corticospinal tract (CST) excitability (Barker, Jalinous, and Freeston 1985), were used to evaluate whether taVNS alters CST excitability independently of voluntary motor behavior.

Together, this multimodal approach - integrating autonomic, ascending neuromodulatory, and cortical measures - offers a robust platform to dissect how taVNS effects are shaped by behavioral state and to uncover neural signatures that reflect its interaction with movement engaged networks. Clarifying these relationships is essential for optimizing stimulation protocols in both basic research and clinical interventions, particularly in neurorehabilitation where taVNS is increasingly paired with motor training.

## METHODS

### Participants

We recruited thirty-six healthy adults to ensure sufficient statistical power. Based on prior effect size estimates, a minimum of 15 participants per measurement modality was required to achieve 80% power at an alpha level of 0.05 (Lerman et al. 2019; Machetanz et al. 2021; Sharon, Fahoum, and Nir 2021). All participants met established inclusion and exclusion criteria for taVNS and TMS (Badran et al. 2019; Rossi et al. 2009). Two participants withdrew after providing informed consent and one was excluded at the start due to inability to feel the stimulation. Twenty-four persons participated in the taVNS-movement physiology experiment (n = 24; 24.2 ± 2.4 years old; 6 females) and fifteen in the taVNS CST excitability experiment (n = 15; 24.8 ± 2.9 years; 4 females), with some overlap between groups. No participants reported any major side effects resulting from the stimulations. The study was approved by the ETH Zurich Ethics Commission (EK 2023-N-316) and was performed in accordance with the Helsinki Declaration of the World Medical Association (World Medical Association 2013).

### taVNS

Stimulation was delivered by an in-house developed current-controlled pulse generator operating on a custom graphical user interface (GUI) written in Python. Electrolyte solution (Signa Spray) was applied to the electrodes. A custom electrode targeting the superior cymba conchae and inferior cavum conchae of the left ear meatus was utilized for taVNS (Fig. 1C). Sham used a modified ear clip (Vagustim^TM^) electrode placed on the left earlobe (Fig. 1C) that has weaker vagal innervation as described previously (Kraus et al. 2013). Bipolar square wave pulses of 25Hz (250us pulse width) 2 second stimulation trains were used in both taVNS and sham conditions, while no stimulation was delivered in the control condition. Each stimulation condition was calibrated separately in increments of 0.1 mA up to the subjective pain threshold and reduced to 90% or up to a maximum of 5mA and 50V (Supplemental Fig. 1A, C). The sham condition allowed control for nonspecific effects of taVNS by mimicking the sensory experience without inducing the associated neural activation (Kraus et al. 2013), ensuring that any observed differences in measures could be attributed specifically to stimulation of the auricular vagus nerve. Importantly, higher currents were required for sham stimulation to reach the same subjective perception as taVNS (Fig. S1).

**Fig. 1.**
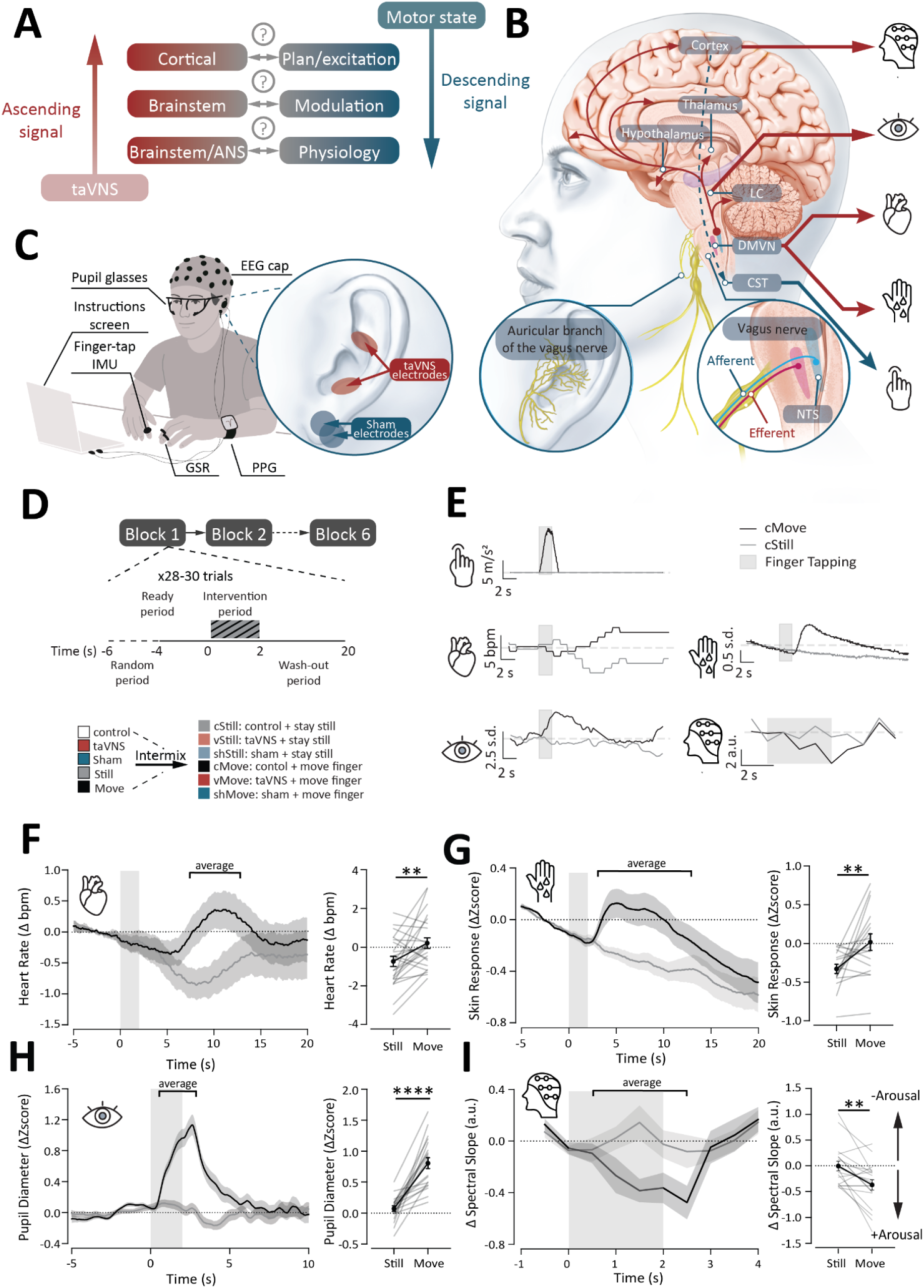
Experimental concept, design and effects of movement alone on the neurophysiological outcomes. A) Conceptual schematic of potential interactions between taVNS signal cascade and motor control. B) Anatomical pathways of ascending taVNS signal (red), descending corticospinal pathway (blue dashed), and associated neurophysiological readouts. LC = locus coeruleus, DMVN = dorsal motor vagus nucleus, CST = corticospinal tract. C) taVNS-Movement Physiology Experiment setup. D) Schematic of the experiment, trial structure and experimental groups. E) Representative traces of single trial data for cStill (grey) and cMove (black) conditions: finger tapping, HR, GSR, pupil diameter, and spectral slope. F-I) Time course plots (left) and selected periods for pairwise analysis (right) for HR, GSR, pupil diameter and whole brain spectral slope respectively. Grey box indicates the two second cue for movement, bar indicates period for averaging and pairwise comparisons. Data are mean +/- SEM. Paired t-test, ** = p<0.01, **** = p<0.0001.

### taVNS-Movement Neurophysiology Experiment

The experimental set-up is shown in Fig. 1C. The experiment was performed in a quiet room with constant lighting where participants were seated in front of a screen with instructions. Electrodes were placed and the stimulation was calibrated. Measurement devices for finger tapping, HR, GSR, pupil diameter and EEG were fitted and the experiment commenced. Stimulation control and data acquisition were synchronised by a master computer. The experiment was split into a maximum of 12 blocks, consisting of 26 to 30 trials each, spread over 2 days. Trial conditions (e.g. move vs. still x control (c) vs. sham (sh) vs. taVNS (v)) were randomized per block. Six conditions (cMove, cStill, shMove, shStill, vMove, vStill) were analysed in the present study. All modalities were recorded continuously throughout the block. Each trial epoch consisted of a ready (randomised between 4 and 6s), intervention (2s) and wash-out (18s) phases. The ready phase was signalled with the word “index” displayed on the screen. An auditory tone (440 Hz; 2s; stereo) cued participants to perform repetitive, fast index finger tapping throughout the tone. The absence of sound cue instructed participants to remain still. The following washout period was intended for the measured parameters to return to baseline. Participants were instructed to remain still and relaxed unless prompted otherwise by the auditory tone (Fig. 1D).

### Finger tapping

Finger movements were captured using an inertial measurement unit (IMU, Movella Dot) placed on the right index finger’s distal interphalangeal (FDI) joint, and wirelessly transmitting live data (60Hz) to the central computer via bluetooth. The acceleration magnitude was calculated by applying the euclidean norm to raw accelerometer measures. Data was epoched to a window of -5 to 20 seconds, where 0 seconds indicates intervention cue (i.e., for movement and/or stimulation start, depending on the condition). Trials with no viable IMU data, movement trials with peak absolute acceleration under 2m/s^2^ during the intervention window (0 to 2 seconds), and still trials with any acceleration greater than 1m/s^2^ were discarded. In total, 345 out of 1877 trials were excluded from further analysis (a total of n = 24 participants data analyzed).

### Heart rate

HR was measured via a wrist-worn photoplethysmogram (PPG) Grove sensor (101020082, Seeed Studio), continuously streaming beat per minute (BPM) data at 20 Hz to the master computer via an Arduino. The unphysiological values for typical resting state HR (<40 or >110 BPM (Bonnemeier et al. 2003)) were converted to Not a Number (NaN) values. The data were epoched, and epochs were discarded if they contained more than 25% invalid data or if there was a change greater than 20 BPM across the trial. Otherwise, data gaps were linearly interpolated, resulting in 1190 analyzed trials (a total of n = 24 participants data analyzed).

### Galvanic skin response

GSR was acquired using the Grove sensor (101020052, Seeed Studio), which was calibrated at the start of the session. Live GSR data was sent via an Arduino to the master computer and was resampled at 20 Hz. Out-of-range values (i.e., flat values, reaching outside of the dynamic sensor range) were manually excluded by conversion to NaN values. Data was epoched to windows of -5 to 20 seconds and smoothed with a moving mean filter of 500ms. To exclude trials exhibiting nonphysiological baseline variability, a linear slope was fitted to the baseline period (−5 to 0 seconds) of each trial. Trials with slope values exceeding ±6 standard deviations (SD) from the participant-specific median were discarded. The epoch-normalized Zscore was then calculated as follows to preserve response amplitude without the tonic (e.g., sweating due to GSR finger sleeves) effects: each epoch value was local-baselined by subtracting the 5s pre-stimulation average, followed by calculating global, post-baseline SD, and dividing each sample by the SD. Epochs with more than 40% of invalid data were discarded. Participants with one or no trials per condition were discarded from analysis, resulting in 626 trials (and a total of n = 17 participants data analyzed).

### Pupil diameter

The pupil diameter was captured at 120Hz with the Pupil Core (Pupil Labs) glasses and Pupil Capture software. The left eye was selected for analysis and the 3D pupil model was used to estimate pupil diameter in mm. Blinks and poor estimates were discarded by converting all <90% confidence data points, as calculated by the inbuilt Pupil Core model, to NaN values, and preprocessing as previously described (Weijs et al. 2025). In brief, pupil-diameter speed values that fell outside the median absolute deviation (MAD; multiplier set to 12, repeated 4 times (Kret and Sjak-Shie 2019)), indicative of abnormal dilation speeds, were discarded. Additionally isolated samples with maximum widths of less than 250 ms bordering a gap larger than 40ms were removed. Data was then smoothed using a zero-phase low-pass filter with cutoff frequency of 4Hz (Kret and Sjak-Shie 2019). Epochs with more than 40% of NaN values were discarded, resulting in 1104 trials analyzed (a total of n = 20 participants data analyzed). Pupil diameter was then epoch-normalized (Zscore) as described for the GSR. Finally, the averaged participant data was smoothed using a moving mean filter of 500ms.

### EEG

EEG data was acquired using ANT Neuro eego mylab with a standard 64-channel 10-20 montage and sampled at 2000 Hz. Analysis was conducted using the MNE-Python library (Gramfort et al. 2013). First, a common-average reference was computed for each session, followed by resampling to 500 Hz and band-pass filtering between 1 and 100 Hz. To address high-amplitude, transient stimulation artifacts, artifact subspace reconstruction (ASR; asrpy package, default parameters, cutoff = 20) was used (Blum et al. 2019). The data was then epoched into -1 to 5 second windows. Non physiological epochs and channels were identified and corrected using the AutoReject algorithm (Jas et al. 2017). A 50 Hz notch filter was used to remove line noise.

Next, independent component analysis was performed to remove blink, muscle, and other non-neural artefacts. Further non physiological trial exclusion was performed (e.g., flat signals within [-0.25 μV, 0.25 μV]), removing technical montage errors (observed in 5 participants), or amplitudes outside the range of [-100 μV, 100 μV], resulting in 1068 analyzed trials (n = 19 participants). Finally, the cleaned data was band-pass filtered between 1 and 45 Hz for further analysis.

For spectral slope analysis, a time-frequency decomposition was performed per trial using Morlet wavelet convolution (MNE-Python library) in 0.25 Hz increments. Time-frequency responses (TFR) were then segmented per channel and per epoch between -1 to 0 and 0 to 4 seconds in increments of 0.5 seconds. All TFR responses were then clipped and linearly interpolated from 22-28Hz to remove any possibility of remaining stimulation artifact using the FOOOF toolbox interpolation method (Donoghue et al. 2020). The aperiodic component of the power spectra was calculated (FOOOF toolbox, default settings (Donoghue et al. 2020)) and the 30-45 Hz frequency range was selected for analysis as a marker of inhibition-excitation balance and arousal in animal and human models (R. Gao, Peterson, and Voytek 2017; Weijs et al. 2025; Lendner et al. 2020). Spectral slopes of the response event (0 to 4 seconds) were then normalized to the baseline window (-1 to 0 seconds). For statistical analyses average spectral slopes between 0.5 to 2.5 seconds were used. For sensorimotor region analysis, CP3, CP5, C3, C5 electrodes were selected as previously reported to map to the regions of the finger (D’Ambrosio et al. 2022). Channels of interest were averaged across trials for statistical analyses.

### taVNS Corticospinal Tract Excitability Experiment

The experiment was intended to probe CST excitability during taVNS and the set-up is shown in Fig. 4A. Although MEPs predominantly reflect cortical output, contributions from spinal excitability cannot be entirely ruled out. Therefore, the term CST is used here to encompass the entire corticospinal pathway. During the measurement, participants sat comfortably at rest and directed their gaze toward a fixation cross. At the beginning of the experiment, participants were equipped with an electromyography (EMG) electrode over their right first dorsal interosseous (FDI) muscle, for MEP of the FDI to be used as a primary physiological read-out. The experiment consisted of three blocks of 15 trials each, with conditions (no stimulation control vs. taVNS vs. sham) randomized per block. Before each block, two TMS pulses separated by 4 seconds were delivered every 6-8 seconds for 10 repetitions, to establish a baseline of the muscle response before applying intermixed stimulation. For sham and taVNS conditions, TMS was delivered 1s into the taVNS train, followed by a second pulse 4 seconds later and a 6-8 second washout period. For control, only two TMS pulses separated by 4 seconds were delivered (Fig. 4F). For each condition, 15 MEPs were recorded. The experimenter holding the TMS coil was blinded to the stimulation condition.

### TMS Properties

Single-pulse monophasic TMS was delivered over the left primary motor cortex using a 70 mm figure-of-eight coil connected to the Magstim 200 stimulator (Magstim, UK). The coil was positioned over the hotspot of the right FDI muscle. The hotspot was defined as the stimulation site where TMS delivery resulted in the most consistent and largest MEPs in the resting muscle. The coil was held tangential to the surface of the scalp, with the handle pointing backward and laterally at 45° away from the nasion-inion mid-sagittal line, resulting in a posterior-anterior direction of current flow in the brain. This orientation is considered most effective for inducing an electric field perpendicular to the central sulcus resulting in the stimulation of primary motor cortex neurons (Rathelot and Strick 2009); (Mills, Boniface, and Schubert 1992). The optimal coil location was registered using a neuronavigation software (Brainsight Frameless, Rogue Research Inc.). The position of the participant’s head and TMS coil was constantly monitored in real-time with the Polaris Vicra Optical Tracking System (Northern Digital Inc.). This ensured that the center of the coil was kept within 2 mm of the determined hotspot, and that the coil orientation was consistent throughout the experiment. The resting motor threshold (RMT) was determined for each participant at the start of the session. RMT was defined as the lowest TMS intensity to elicit MEPs with peak-to-peak amplitude ≥ 0.05 mV in the relaxed muscle, in five out of 10 consecutive trials, i.e., 50% ((Rossini et al. 1994); mean RMT = 43.26 ± 5.04% of the maximum stimulator output, MSO, range: 33-52%). Single-pulse TMS was delivered at an intensity of 120% of the individual RMT during the experiment (52.2 ± 5.97% MSO, range: 40–63%).

### EMG and data analysis

The muscle response was recorded by a wireless surface EMG electrode (Trigno Wireless, Delsys) placed over the right FDI muscle. Raw signals were amplified (sampling rate, 2 kHz), digitized with a CED micro 1401 AD converter and Signal software V2.13 (Cambridge Electronic Design), and stored for off-line analysis. Timing of the TMS delivery and EMG data recording were synchronized by a Python script connected to the CED via a custom microcontroller. Muscular relaxation was constantly monitored through visual feedback of EMG activity and participants were instructed to relax their muscles if necessary.

Pre-processing of the EMG data was conducted using MATLAB R2020a (MathWorks, Inc., Natick, MA, USA). The EMG data was band-pass filtered (30-800 Hz). Filtering was applied separately for the pre-TMS background EMG (bgEMG; measured for 100 ms between 105 ms and 5 ms before the TMS pulse) and post-TMS period containing peak-to-peak MEP amplitude to avoid smearing the MEP into bgEMG data. The data was filtered with a high-pass, 30 Hz, 2nd-order butterworth filter and subsequently a low pass, 800 Hz, 2nd order butterworth filter. Additionally, a 50Hz IIR notch filter (Q=25, bandwidth=2Hz) was applied to the bgEMG signal only. Peak-to-peak amplitude was defined as the maximum voltage difference within 15 to 60 ms after the TMS pulse. Trials were excluded from analysis if the bgEMG activity during the pre-stimulation period deviated by more than 2 standard deviations from the participant’s mean bgEMG across all trials, resulting in 94 discarded trials of 2250 total trials (a total of n = 15 participants data analyzed). MEP amplitudes from each stimulation condition were normalized to the mean baseline values recorded at the beginning of each corresponding block. The normalized data were expressed as percentage change from baseline.

### Statistical analysis

For both experiments, epoch data were averaged per participant and exported to Prism 10.4.1 (GraphPad) or python scipy stats toolbox for statistical analysis. We first examined the effects of movement on each measure, identifying the specific time points at which movement alone significantly modulated the signal. Multiple t-tests were performed and corrected with Šídák’s test to determine specific time points at which cMove significantly differed from cStill (p<0.05) and were rounded to the nearest 0.5 seconds (indicated as “average” bar in Fig. 1F-I). Where indicated, these timepoints were used for subsequent analyses of stimulation-related effects. To evaluate differences between stimulation conditions within each behavioral context, we conducted paired t-tests across condition pairs with appropriate correction for multiple comparisons. For analyses involving six pairwise comparisons (cStill vs. vStill, vStill vs. shStill, cStill vs. shStill, cMove vs. vMove, vMove vs. shMove, cMove vs. shMove; Fig. 2-4), p-values were adjusted using the Benjamini-Hochberg false discovery rate procedure. For analyses involving only three comparisons (e.g., Fig. 4B,C), Holm-Sidak corrections were applied. Data are reported as mean ± SEM unless stated otherwise. Statistical significance was defined as p < 0.05 and presented p-values are all corrected.

**Fig. 2.**
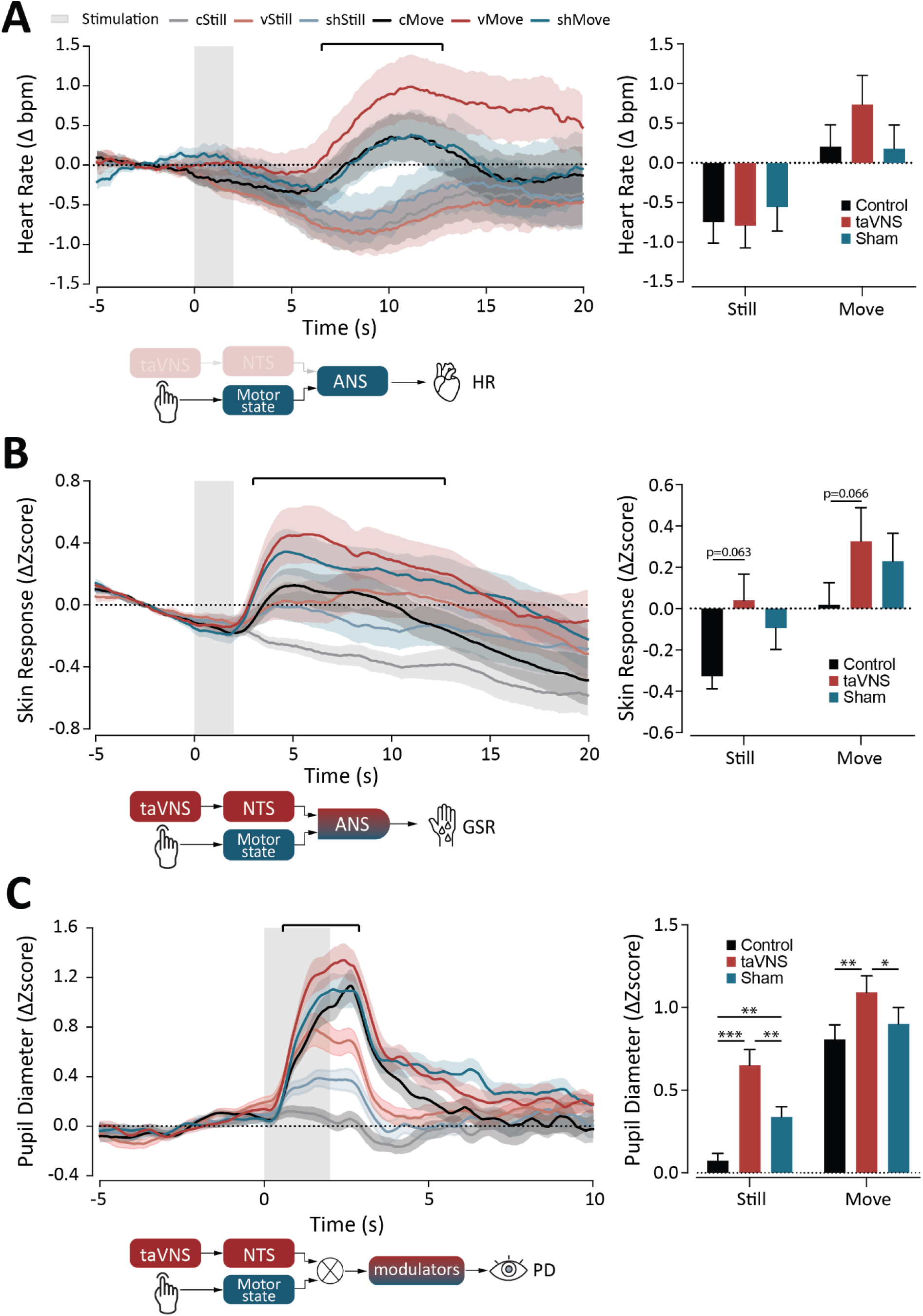
Divergent recruitment of autonomic and ascending neuromodulator associated measures by taVNS and movement. A) Time-course (left) of the epoched heart rate (HR) changes (Δ bpm) and selected average periods used for analysis (right). Grey box indicates the two second cue for movement and stimulation, bar indicates period for averaging and statistical analysis (same as in Fig. 1). Data are mean +/- SEM. B) Same as in A for GSR (Δ Zscore). C) As in A and B for changes in pupil dilation (Δ Zscore). * = p<0.05, ** = p<0.01, *** = p<0.001.

## RESULTS

Conceptually, taVNS triggers an ascending signal cascade, propagating through brainstem autonomic centers and neuromodulatory nuclei, and eventually reaching cortical regions. Concurrently, the motor control hierarchy progresses from contextual planning to execution, while integrating with physiological states via the ANS (Fig. 1A). Given the functional and anatomical overlap, we sought to determine whether the taVNS had effects on neurophysiological markers associated with different underlying neural circuits during a movement task (Fig. 1B, C). We employed a multi-modal experimental setup (Fig. 1C, D) to measure the effects of taVNS on motor and physiological responses across a cohort of healthy participants.

### Finger tapping is strongly represented in autonomic, ascending neuromodulatory, and cortical activity measures

All of the employed neurophysiological measures showed significant differences between cMove and cStill conditions. Activation of ANS activity markers, the HR and GSR, was significantly higher in cMove compared to cStill (Fig. 1F, G, HR: t(23) = 3.13, p = 0.0047; GSR: t(16) = -3.058, p = 0.0075). Pupil dilation and brain-wide spectral slope (all channels), markers of the neuromodulatory system and arousal, respectively, were significantly elevated in cMove compared to cStill (Fig. 1H, I, pupil diameter: t(19) = -7.504, p < 0.0001; spectral slope: t(18) = 3.711, p = 0.0016). These results confirm that movement robustly modulates autonomic, ascending neuromodulatory and arousal markers, emphasizing the importance of accounting for motor activity when evaluating taVNS effects.

### taVNS activated neuromodulatory markers with limited effects on ANS markers

taVNS was investigated for its effects on autonomic and neuromodulatory markers, revealing distinct responses in HR, GSR and pupil diameter across still and move conditions. No significant effects of taVNS was detected in HR (Fig. 2A, cStill x vStill: t(23) = 0.133, p = 0.944; vStill x shStill: t(23) = -0.642, p =0.839; cStill x shStill: t(23) = -0.593, p = 0.839; cMove x vMove: t(23) = -1.176, p = 0.755; vMove vs shMove: t(23) = 1.324, p = 0.755; cMove x shMove: t(23) = 0.071, p = 0.944), nor in GSR measures, the latter showed a non-significant trend toward greater activation in the taVNS condition compared to control and sham in both movement paradigms (Fig. 2B, cStill x vStill: t(16) = -2.899, p = 0.063; vStill x shStill: t(16) = 0.907, p = 0.453 cStill x shStill: t(16) = -2.112, p = 0.101; cMove x vMove: t(16) = -2.535, p = 0.066; vMove vs shMove: t(16) = 0.631, p = 0.537; cMove x shMove: t(16) = -1.933, p = 0.107). In contrast, the pupil dilation response during taVNS was significantly greater compared to both sham and control across both still and move conditions (Fig. 2C, cStill x vStill: t(19) = -5.191, p = 0.0002; vStill x shStill: t(19) = 3.061, p = 0.013; cStill x shStill: t(19) = -3.318, p = 0.007; cMove x vMove: t(19) = -3.690, p = 0.005; vMove vs shMove: t(19) = 2.634, p = 0.020; cMove x shMove: t(19) = -1.454, p = 0.162). These findings suggest that transient taVNS selectively engages neuromodulatory pathways regardless of movement, while its effects on autonomic markers remains limited.

### Cortical effects show regional specificity rather than global arousal

Global cortical arousal, which would manifest as a widespread increase in spectral slope across all electrode sites ((Weijs et al. 2025); Fig. 3A), was not significantly influenced by stimulation in either still or movement (Fig 3B, cStill x vStill: t(18) = 1.183, p = 0.464; vStill x shStill: t(18) = -0.133, p = 0.896; cStill x shStill: t(18) = 1.046, p = 0.464; cMove x vMove: t(18) = 2.033, p = 0.189; vMove vs shMove: t(18) = -1.981, p = 0.189; cMove x shMove: t(18) = -0.429, p = 0.808). However, examination of electrodes over the sensorimotor cortex, selected based on their anatomical correspondence to hand sensorimotor representation areas (Fig. 3A, (D’Ambrosio et al. 2022)), revealed a significantly flatter slope during vMove compared to cMove and shMove (Fig. 3C, cStill x vStill: t(18) = 1.330, p = 0.300; vStill x shStill: t(18) = 0.867, p = 0.397; cStill x shStill: t(18) = 2.366, p = 0.059; cMove x vMove: t(18) = 2.735, p = 0.041; vMove vs shMove: t(18) = -3.194, p = 0.030; cMove x shMove: t(18) = -1.116, p = 0.335). Thus, taVNS was associated with increased spectral slope in the sensorimotor region only during movement, with no significant effect during Still. This pattern of modulation appears to be gated by the movement behaviour and is unlikely to reflect a sensory perception artifact due to lack of detected effect of sham stimulation.

**Fig. 3.**
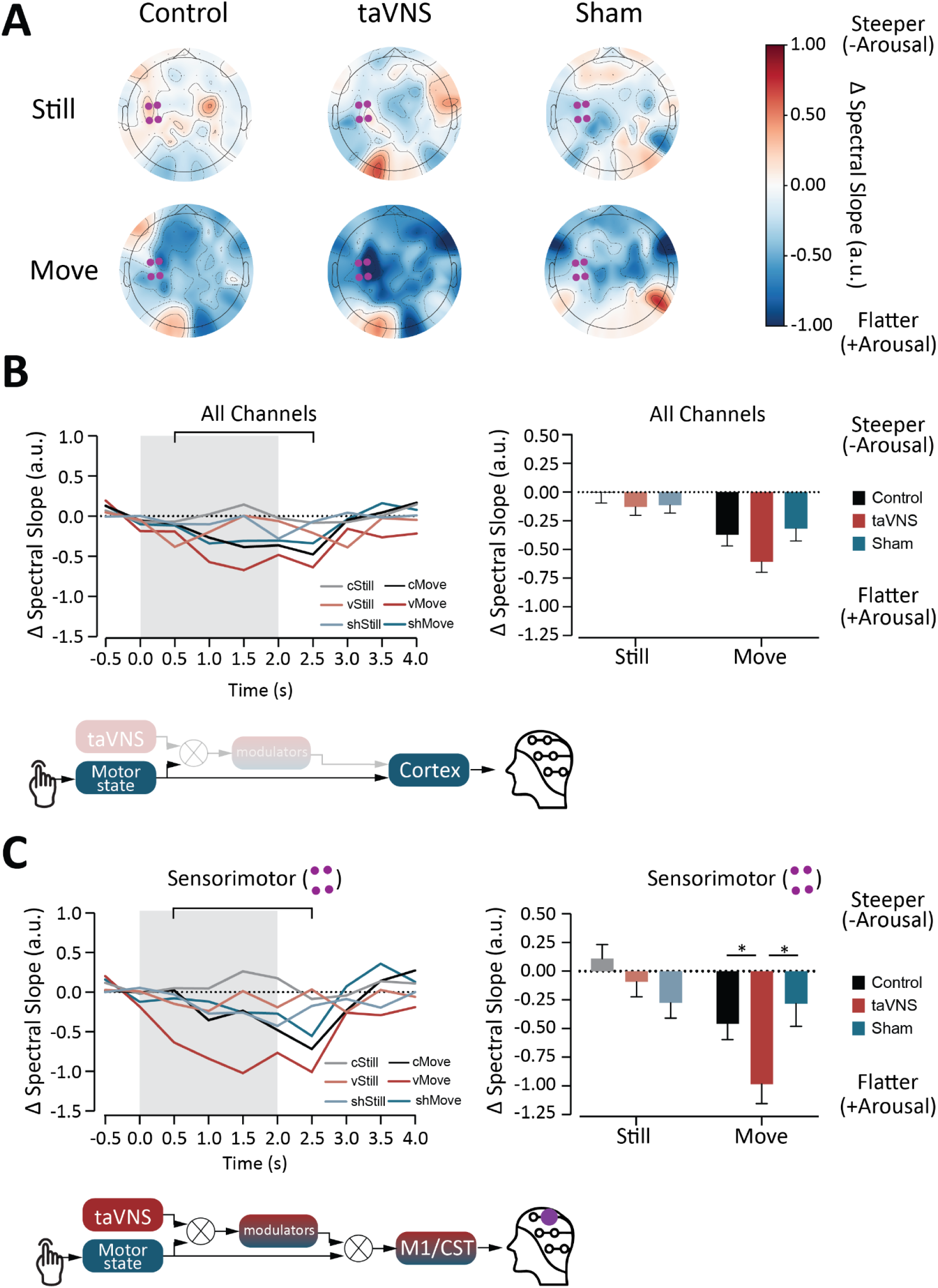
Sensorimotor cortex-localized taVNS effects on EEG spectral slope contrast with global cortical activation during movement A) Topographic plots of EEG-measured average change in epoched spectral slope (0.5 to 2.5 seconds). Cooler colors (blue) indicate flatter slopes associated with increased arousal, while warmer colors (red) indicate steeper slopes associated with reduced arousal. Purple dots mark electrodes over the sensorimotor cortex (n = 19). B) Time-course (left) of the epoched Δ spectral slope across all channels and selected average periods used for analysis (right). Grey box indicates the two second cue for movement and stimulation, bar indicates period for averaging and statistical analysis. C) Same as in B, but only for the selected channels over the left sensorimotor cortex (‘C3’, ‘C5’, ‘CP3’, ‘CP5’ ; indicated in A), * = p<0.05.

**Fig. 4.**
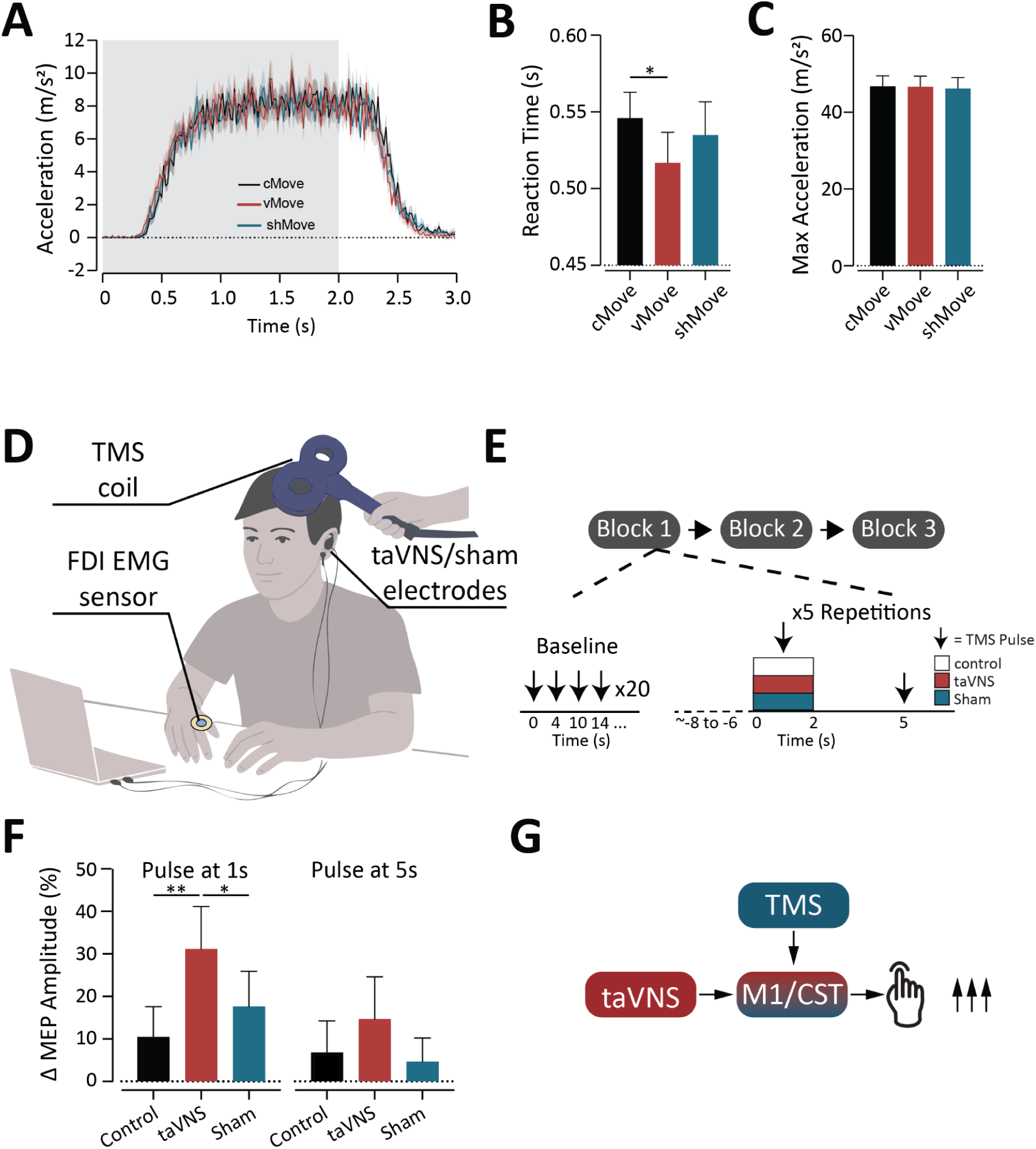
taVNS enhances MEP responses, consistent with transient increase in corticospinal excitability A) Time course of finger tapping acceleration data (m/s^2^). B) Average reaction time (s) to start finger tapping from cue. Reaction time was found for the first time the acceleration crosses 0.5 mm/s^2^. *p<0.05. C) Same as in B, for max acceleration (m/s^2^) during finger tapping. D) taVNS Corticospinal Tract Excitability Experiment setup. E) Schematic of the experimental protocol and trial structure. F) Average ΔMEP amplitude (% change from mean MEP baseline amplitude) during stimulation (at 1s, left), and 4 seconds afterwards (right). * = p<0.05, ** = p<0.01. G) Schematic summary of results. taVNS transiently increases CST excitability, leading to enhanced MEP responses.

### taVNS transiently modulates motor response and enhances CST excitability

We next looked at the behavioral output from the finger tapping task by looking at the acceleration values during the movement trials (Fig. 4A). We found there was a significant decrease in reaction time for vMove compared to cMove but not to shMove (Fig. 4B, cMove x vMove: t(23) = 2.812, p = 0.029; vMove x shMove: t(23) = -1.697, p = 0.196, cMove x shMove: t(23) = 0.970, p = 0.342; Supplemental Fig 2.). However, while taVNS significantly reduced reaction time, the magnitude of finger tapping acceleration did not differ significantly across the three conditions (Fig. 4C, cMove x vMove: t(23) = 0.175, p = 0.863; vMove x shMove: t(23) = 0.735, p = 0.851; cMove x shMove: t(23) = 0.721, p = 0.851).

To assess whether cortical spectral slope changes correspond with variations in CST excitability, independent of behavioral execution, we employed a TMS paradigm where the first pulse was delivered within the taVNS stimulation train and the second pulse after stimulation offset (Fig. 4D, E). Analysis revealed that taVNS produces significantly enhanced MEP amplitudes compared to sham and control conditions when the TMS pulse is delivered during the stimulation period, yet this effect did not persist by the time of the second pulse (Fig. 4F, pulses at 1s: Control x taVNS t(14) = -3.912, p = 0.009; taVNS x Sham t(14) = 2.793, p = 0.043; Control x Sham t(14) = -1.491, p = 0.190; pulses at 5s Control x taVNS t(14) = -1.671, p = 0.175; taVNS x Sham t(14) = 1.803, p = 0.175; Control x Sham t(14) = 0.437, p = 0.669). taVNS transiently reduces motor reaction time and boosts CST excitability, but only during stimulation, suggesting a selective, short-lived modulatory effect.

## DISCUSSION

This study aimed to investigate if transient taVNS during voluntary movement modulates neural and physiological activity and whether there are any movement-taVNS interactions in these neurophysiological measures. First, taVNS effects were analysed in two behavioural states: an active (go) condition involving voluntary finger movement, and a motor inhibition (no-go) condition. We did not detect taVNS effects on HR nor on GSR measures, although there was a non-significant trend for greater responses in GSR in taVNS conditions. We did find significant taVNS effects on pupil diameter during both behavioral states, and observed a movement-gated taVNS activation of the sensorimotor cortex. This taVNS increased CST excitability was confirmed with the TMS experiment.

Together, these results address the critical and largely unaddressed gap of how taVNS interacts with ongoing motor-related neural activity, suggesting that taVNS actively engages and modulates sensorimotor circuits in a behaviorally dependent manner. These findings highlight that the neural impact of taVNS is not static, but shaped by behavioral context, underscoring the importance of aligning taVNS with specific tasks to enhance its impact, and potentially guiding its application in neurorehabilitation protocols.

### Distinct taVNS effects on autonomic and arousal metrics

Numerous studies have reported that taVNS can influence cardiovascular (HR and heart rate variability) and GSR markers, typically interpreted as evidence of parasympathetic engagement or broader autonomic modulation (Geng, Yang, et al. 2022; Badran et al. 2018; G. Tan et al. 2024). Interestingly, our findings reveal a more nuanced pattern in the context of taVNS-movement interactions. While movement robustly modulated both HR and GSR (Fig. 1F,G), there was no observed taVNS effect on autonomic measures, with only a slight trend towards increased GSR activity in taVNS compared to control conditions (Fig 2B). This pattern suggests that the effects of taVNS on ANS may be potentially masked by movement-related autonomic activation. Alternatively, such limited responsiveness could stem from the brief 2-second stimulation epochs used here, in contrast to longer stimulation protocols (e.g., 5 minutes; (Geng, Liu, et al. 2022)) or to paradigms employing physiological gating, such as HR phase-synchronized (Tischer, Szeles, and Kaniusas 2025) or respiration-mediated taVNS (Garcia et al. 2022), both of which have been shown to promote ANS responses.

In addition, taVNS has been associated with shifts in central arousal states, reflected in pupil diameter changes (Skora, Marzecová, and Jocham 2024; Ludwig et al. 2024). In our study, while taVNS had limited effects on traditional autonomic markers, we observed significant taVNS effects on pupil dilation against control and sham conditions in both behaviour paradigms (Fig. 2C), indicating a more robust engagement of neuromodulatory central arousal circuitry. The observed pupil effects likely reflect the recruitment of central neuromodulatory systems that are tightly linked to pupil dynamics, such as the LC norepinephrine, the basal forebrain (BF) cholinergic and LH orexinergic neuropopulations (Viglione, Mazziotti, and Pizzorusso 2023; Grujic et al. 2023; Pfeffer et al. 2022). These systems broadly influence cortical and subcortical networks, contributing to functions such as arousal (Berridge 2008; Villano et al. 2017), attention (Zhang et al. 2023; Maness et al. 2022), neuroplasticity (O’Donnell et al. 2012; Ramanathan, Tuszynski, and Conner 2009; X.-B. Gao and Hermes 2015), and autonomic regulation (Samuels and Szabadi 2008; Berntson, Sarter, and Cacioppo 1998; Nattie and Li 2012). The presence of pupil modulation in the absence of significant HR or GSR changes supports the possibility of a functional decoupling between autonomic and ascending neuromodulatory responses in response to brief stimulation. These findings may suggest that, in our paradigm, phasic taVNS may influence central arousal circuits directly, with limited engagement of downstream autonomic reflex loops such as those mediated by the hypothalamic-pituitary axis or brainstem nuclei like the DMVN.

To probe central arousal further, we assessed EEG-based measures of arousal, focusing specifically on the spectral slope, a marker increasingly recognized for its sensitivity to shifts in cortical excitability and brain state (Weijs et al. 2025; R. Gao, Peterson, and Voytek 2017). Our results demonstrate that while movement significantly affected whole-brain spectral slope, taVNS did not, suggesting that in this paradigm, stimulation did not induce a general cortical arousal response above sham or control conditions (Fig. 3B). This surprising dissociation of pupil diameter and cortical markers of arousal reinforces the likelihood that transient taVNS does not act as a general arousal signal. This in turn raises the question of whether taVNS is selectively modulating a behaviour-dependent neurocircuitry.

### Motor state-specific arousal by taVNS

Given the paradigm’s motor demand, we next investigated whether taVNS specifically modulates EEG activity in sensorimotor regions associated with finger movement. We focused on the EEG spectral slope at electrodes overlying finger motor areas, defined via TMS-evoked MEP mapping (D’Ambrosio et al. 2022). The spectral slope of the sensorimotor-associated electrodes revealed significant effects of taVNS in the movement condition only (Fig. 3C). This pattern suggests a subthreshold shift towards excitation and/or reduced inhibition that is selectively gated by motor behavior. In turn, this results in a spatially restricted modulation contrasting with the absence of global cortical changes - indicating that taVNS does not induce a generalized arousal response.

To assess whether this localized cortical modulation corresponded with changes in motor behavior, we examined reaction times to the go cue. Participants responded significantly faster during vMove compared to cMove (Fig. 4B, Supplemental Fig. 2D). The absence of reaction time differences between sham and control conditions, along with matched sensory intensity ratings across taVNS and sham (Supplemental Fig. 1B), argues against a nonspecific sensory artifact explanation. However, improved response speed was accompanied by a selective increase in false-positive responses during vStill (no-go) trials (Supplemental Fig. 2E), suggestive of reduced inhibitory control. These findings raise the possibility that taVNS amplifies neural sensitivity to stimuli, facilitating rapid, goal-directed actions but simultaneously increasing susceptibility to impulsive responses.

While these behavioral results point to enhanced motor readiness and arousal with taVNS, the reliance on cued, voluntary finger movements limits our ability to disentangle whether these effects stem from movement intention, motor planning, or execution. To address this, we designed a TMS-induced MEP experiment to directly evaluate the excitability of the corticomotor and CST descending pathway, while minimizing the influence of conscious finger motion. We observed significant transient increase in MEP amplitude during taVNS compared to both sham and control conditions (Fig. 4F). This enhancement was not evident in the post-stimulation period, suggesting that taVNS-related increases in CST excitability may be transient. This pattern is broadly consistent with the EEG findings, where sensorimotor spectral slope returned to baseline shortly after stimulation and movement ceased (Fig. 3C).

### Proposed mechanism of action of transient taVNS

Our findings support a model in which transient taVNS functions as a state-dependent modulator of subcortical and cortical excitability, rather than a driver of brain-wide arousal. Instead of inducing overt shifts in autonomic tone, taVNS may subtly alter baseline neural responsiveness, amplifying the impact of ongoing activity in behaviorally engaged circuits. taVNS appears to enhance excitability in sensorimotor regions engaged by the task, as evidenced by spectral slope modulation and TMS-induced MEPs. A plausible underlying mechanism is LC-driven norepinephrine release, which has been shown to be elicited by taVNS in prior fMRI studies (Borgmann et al. 2021; Huang et al. 2023). In turn, norepinephrine was demonstrated to enhance cortical excitability of task-relevant motor circuits, and facilitates rapid, goal-directed actions by amplifying motor-related gain and suppressing background noise (Bouret and Sara 2005; McGinley, David, and McCormick 2015). In terms of the Adaptive Gain framework, our data fit the proposed role of phasic LC signaling promoting an increased attention and a bias towards exploitation (Aston-Jones and Cohen 2005). Note that a recent study showed that 4 seconds taVNS also affected pupil diameter and increased accuracy of perceptual decision-making task without changing the reaction time (Su et al. 2025). Future studies will be required to determine whether the observed effects stem primarily from the more tonic stimulation protocol (4 seconds vs. 2 seconds) or from variations in the experimental task. Additional downstream taVNS-effect contributions may come from BF cholinergic neurons, which modulate motor cortical excitability and attentional gating (Li and Hollis 2021; Kuo et al. 2007) and enhance signal-to-noise ratios in sensorimotor regions (Sarter et al. 2005). Moreover, LH orexin neurons further promote arousal-motor coupling by activating both LC and BF systems and increasing cortical acetylcholine levels, thereby supporting behavioral state transitions that favor movement initiation and sustained task engagement (Fadel and Burk 2010; España et al. 2005; Sakurai 2007). Together, these systems form a distributed gain-control network that taVNS may transiently engage to selectively heighten motor system excitability.

### Impact to taVNS-based interventions

These findings have potential implications for neurorehabilitation, particularly in conditions such as stroke and spinal cord injury, where there is increasing interest in vagus nerve stimulation to enhance motor recovery (Kilgard et al. 2025; Badran et al. 2023; Dawson et al. 2016). In these contexts, enhancing the responsiveness of spared motor circuits is a key therapeutic goal (Campos et al. 2023; Ward and Cohen 2004). The observed context-sensitive modulation suggests that taVNS could serve as a timing-sensitive amplifier of motor system excitability when paired with active movement, optimizing the neural conditions that support motor recovery. This aligns with recent clinical evidence showing that movement-paired taVNS improves motor outcomes more effectively than stimulation delivered independently of behavior (Badran et al. 2023), as seen with invasive VNS (Dawson et al. 2021). While our study focused on transient effects of taVNS, this form of context-contingent engagement of neuromodulatory systems may promote plasticity when paired with repeated training. Future studies will be required to verify taVNS effects in subjects more closely resembling clinical populations as well as to investigate stimulation frequency, intensity and timing relationships, and ultimately implement single-trial multimodal modeling to determine personalized taVNS biomarkers.

## Conclusion

In summary, our findings demonstrate that transient taVNS modulates neural excitability in a behaviorally contingent manner, selectively enhancing activity in sensorimotor circuits during voluntary movement without eliciting broad autonomic or global arousal effects. This context-sensitive neuromodulation appears to engage central arousal systems to amplify task-relevant neural activity, rather than inducing generalized shifts in brain state. These insights not only advance our mechanistic understanding of taVNS but also support its potential as a targeted, movement-paired intervention to enhance motor system responsiveness, laying the groundwork for future applications in neurorehabilitation.

## CRediT authorship contribution statement

1. **C. Perrin:** conceptualisation, methodology, software, formal analysis, investigation, data curation, writing - original draft, writing - review & editing.
2. **F. Pallotti:** conceptualisation, methodology, software, formal analysis, investigation, data curation, writing - original draft, writing - review & editing.
3. **T. Weilenmann:** conceptualisation, methodology, software, investigation, data curation.
4. **C. Lhoste:** writing - original draft, writing - review & editing.
5. **W. Potok-Szybinska:** methodology, writing - review & editing.
6. **X. Zhang:** methodology, writing - review & editing.
7. **N. Wenderoth:** resources, writing - review & editing.
8. **O. Lambercy:** conceptualisation, funding acquisition, resources, writing - review & editing.
9. **D. Donegan:** conceptualisation, funding acquisition, formal analysis, writing - original draft, writing - review & editing.
10. **P. Viskaitis:** conceptualisation, funding acquisition, writing - original draft, writing - review & editing.

## Acknowledgments

The authors thank the participants for their time and contribution to this work. Additionally they would like to thank Andrea Anliker, Abigail Cécile Vogel, Ladina Flavia Wohlwend, David Mijajlovic, Manuel Glahn, Léa Pistorius, Marine Bruttin, and Nick Lenzin for their contributions to the development of the device, data processing and acquisition. The authors are also grateful to Prof. Denis Burdakov and Prof. Roger Gassert for their support of the project and review of the manuscript.

---

**Supplementary Fig. S1.**
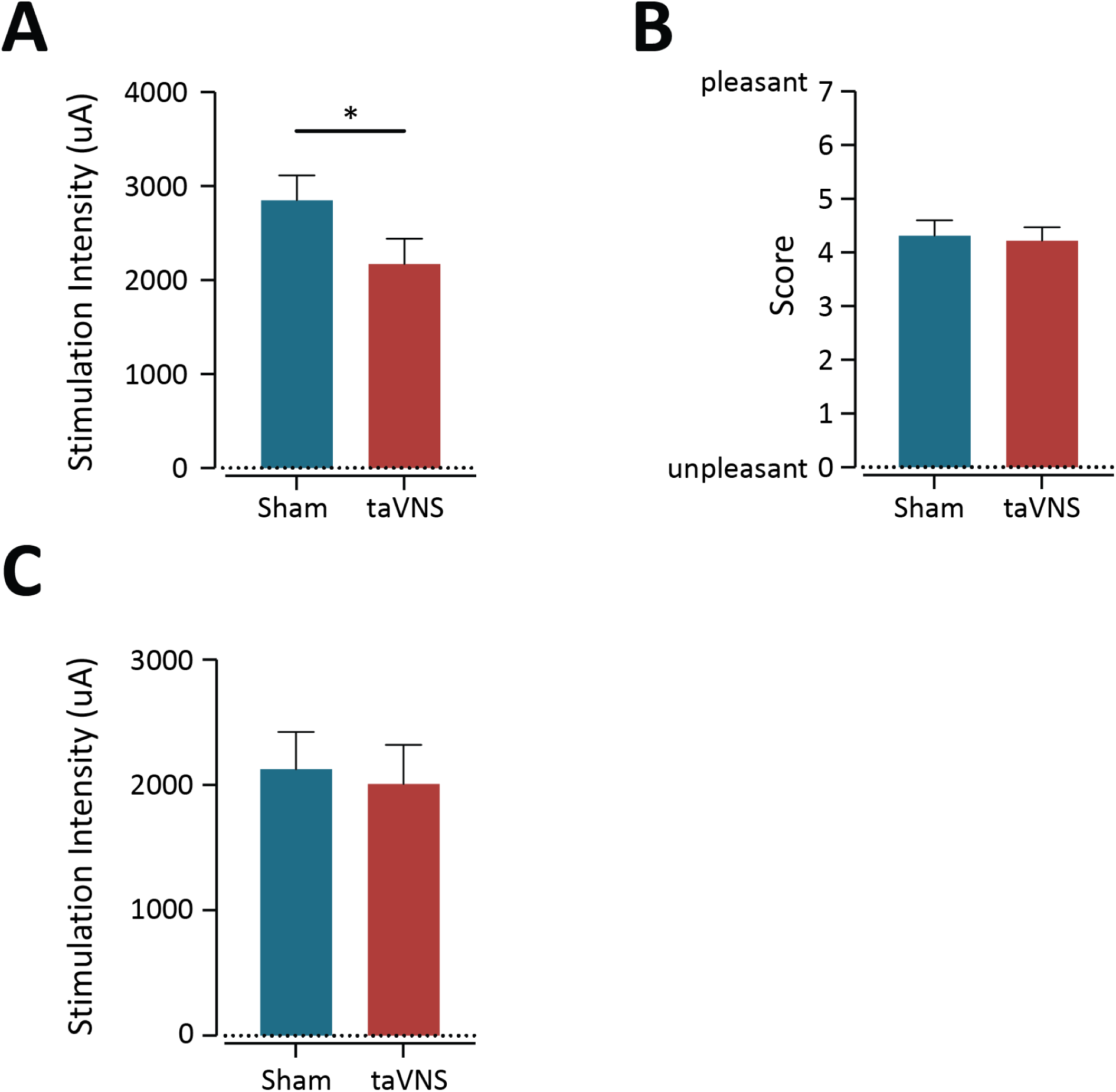
A) Average stimulation intensity calibrated for taVNS-Movement Physiology Experiment. Data shown are mean +/- SEM. Paired t-test (t(23) = 2.359, p = 0.0272). B) Questionnaire score for pleasantness of sham and taVNS stimulation. To the question “I found stimulation pleasant and enjoyable”, scores are out of 7, with 1 = unpleasant and 7 = very pleasant. Data shown are mean +/- SEM. Paired t-test (t(23) = 0.4987, p = 0.6229). C) Average stimulation intensity calibrated for the taVNS Corticospinal Tract Excitability Experiment. Data shown are mean +/- SEM. Paired t-test (t(14) = 0.4458, p = 0.6626)

**Supplementary Fig. S2.**
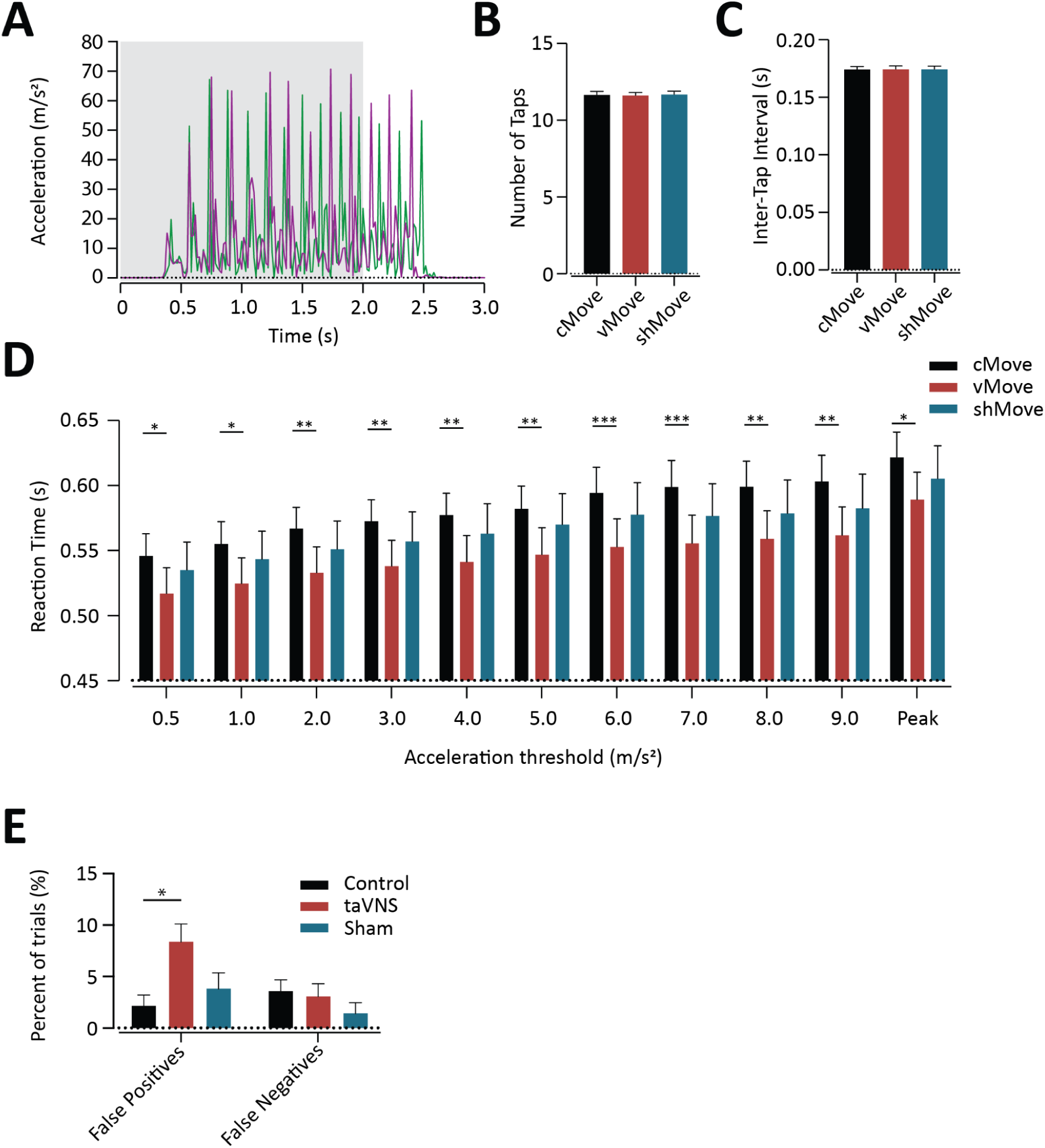
A) Two example raw acceleration traces aligned to movement cue (grey bar). B) Average number of finger taps during the movement cue period. Data shown are mean+/- SEM. Paired t-tests: cMove x vMove: t(23) = 0.364, p = 0.921; vMove x shMove: t(23) = -0.856, p =0.784 ; cMove x shMove: t(23) = -0.257, p = 0.921. C) Average interval between finger taps during the movement period. Data shown are mean +/- SEM. Paired t-tests: cMove x vMove: t(23) = -0.122, p = 0.997; vMove x shMove: t(23) = -0.058, p = 0.997; cMove x shMove: t(23) = -0.180, p = 0.997. D) Average reaction time (s) to start finger tapping from cue, with reaction time found as the first time the acceleration crosses different thresholds (m/s^2^, from 0.5 to the time of the first acceleration peak). REML 2 way anova: Significant main effects of acceleration threshold (F(2.156, 49.59) = 40.36, p<0.0001) and stimulation (F(1.852,42.59) = 5.660, p = 0.0078) were detected, without interaction (F(4.063,93.45) = 0.8522, p = 0.4973). Errorbars indicate Tukey’s multiple comparisons across the different time points. * = p<0.05, ** = p<0.01, *** = p<0.001 E) Percent of total trials for each condition considered as false positives (fp, i.e., still trials with any acceleration greater than 1 m/s^2^ during the intervention window (0 to 2 seconds)) or false negatives (fn, i.e., movement trials with peak absolute acceleration under 6m/s^2^ during the intervention window). Paired t-tests cStill fp x vStill fp: t(23) = -2.983, p = 0.040; vStill fp x shStill fp: t(23) = 2.046, p =0.105; cStill fp x shStill fp: t(23) = -0.798, p = 0.520; cMove fn x vMove fn: t(23) = 0.372, p = 0.713; vMove fn x shMove fn: t(23) = 2.191, p =1.05; cMove fn x shMove fn: t(23) = 1.798, p = 0.128.

